# Type III interferon signaling restricts Enterovirus 71 infection of goblet cells

**DOI:** 10.1101/344531

**Authors:** Charles Good, Alexandra I. Wells, Carolyn B. Coyne

**Author notes:** Address correspondence: Carolyn Coyne, PhD, 9116 Rangos Research Center, Children’s Hospital of Pittsburgh of UPMC, One Children’s Hospital Way, 4401 Penn Avenue, Pittsburgh, PA 15224.

## Abstract

Recent worldwide outbreaks of enterovirus (EV71) have caused major epidemics of hand, foot, and mouth disease (HFMD) with severe neurological complications, including acute flaccid paralysis. EV71 is transmitted by the enteral route, but very little is known about the mechanisms it utilizes to cross the human gastrointestinal (GI) tract. Using primary human intestinal epithelial monolayers, we show that EV71 infects the GI epithelium from the apical surface, where it preferentially infects goblet cells. Unlike echovirus 11 (E11), an enterovirus that infects enterocytes, EV71 infection did not alter epithelial barrier function, but did reduce the expression of a goblet cell-derived mucin, suggesting it alters goblet cell function. We also show that the intestinal epithelium responds to EV71 infection through the selective induction of type III IFNs, which potently restrict EV71 replication. Collectively, these findings define the early events associated with EV71 infections of the human intestinal epithelium and show that host IFN signaling controls replication in an IFN-specific manner.

## Introduction

Enteroviruses are small (∼30nm) single stranded RNA viruses that cause a broad spectrum of illness in humans. Disease manifestations of enterovirus infections can range from acute, self-limited febrile illness to meningitis, endocarditis, acute paralysis, and even death. Enterovirus 71 (EV71) has been associated with major epidemics of hand, foot, and mouth disease (HFMD) worldwide and severe neurological complications, including meningitis, encephalitis, and acute flaccid paralysis^1^. First identified in 1969^2^, EV71 outbreaks have occurred throughout the globe, with epidemics most commonly occurring in the Asia-Pacific region. Between 2008-2012, outbreaks of EV71 in China have been associated with over 7,000,000 cases of HFMD and almost 2500 deaths^3^. The pediatric population is at greatest risk for developing EV71-associated complications, with the vast majority of fatalities occurring in children below the age of two^3^^−^^6^. There are currently no approved therapeutics to treat or prevent EV71 infections.

EV71 is transmitted by the fecal-oral route, where it targets the human gastrointestinal (GI) epithelium for host invasion. The mechanisms utilized by EV71 to cross the GI epithelial barrier remain largely unknown, owing in part to the lack of *in vivo* models to study EV71 infections by the enteral route. For example, modeling EV71 infections in mouse models is complex given the need to rely on the use of mouse adapted viral strains, animals lacking functional interferon (IFN) signaling, and/or mice overexpressing the human homolog of the primary EV71 receptor SCARB2^7^^−^^12^. Previous work in non-human primate models parallel the CNS complication associated with EV71 infections in humans, including when infected by the enteral route^9, 13, 14^. However, despite the development of these models, which provide platforms to determine the efficacy of EV71 vaccines and therapeutics in animals, the specific mechanisms by which EV71 crosses the human GI epithelial barrier have yet to be defined.

The human GI epithelium is a complex cellular barrier composed of multiple cell types, including those of absorptive (enterocytes) and secretory (goblet, enteroendocrine, and Paneth) lineages. These diverse cell types are derived from Lgr5^+^ stem cells located within the base of intestinal crypts, which differentiate into absorptive and secretory lineages^15^. Major advances in the development of *ex vivo* ‘mini-gut’ enteroid models, in which primary human intestinal crypts are isolated and cultured into epithelial structures that differentiate to contain the multiple cell types present in the human intestine^16^^−^^18^, have significantly expanded our understanding of enteric virus-GI interactions (reviewed in ^19^). In previous work, we utilized enteroids isolated from human fetal small intestines to profile the susceptibility of the human intestine to enterovirus infections, using echovirus 11 (E11), coxsackievirus B (CVB) and EV71 as models^20^. We showed that E11 exhibits a cell type specificity of infection and infects both enterocytes and enteroendocrine cells but is unable to infect goblet cells^20^, suggesting that enteroviruses exhibit a cell type specificity in their infections of the GI epithelium. However, during these studies, we noted that in contrast to both E11 and CVB, EV71 replicated to low levels in human enteroids, although the mechanistic basis for this remained unclear^20^.

Although the crypt-based model utilized in our previous work has many advantages over standard cell line-based models, the culturing of crypts in Matrigel induces the formation of 3-D structures wherein the luminal (apical) domain faces inward and the basolateral domain faces the culture medium. This impacts the polarity by which viruses infect enteroids, restricts the ability to determine whether there is a polarity of viral entry and/or release, and precludes an assessment of alterations that may be induced to the epithelium by infection, such as loss of barrier function. Here, we developed a monolayer model using isolated human fetal crypts cultured on permeable porous membrane inserts, which leads to the formation of a single cell monolayer containing all of the distinct cell types present in the GI epithelium. Using this model, we found that E11 and EV71 exhibit differences in their ability to bind and infect from the apical or basolateral surfaces, with a strong basolateral polarity for E11 and an apical polarity for EV71. Interestingly, we found that whereas E11 targets enterocytes and abolishes epithelial structure and barrier function, EV71 preferentially infects goblet cells and infection reduces the expression of a goblet cell-derived mucin. Lastly, we show that EV71 infection specifically induces the type III IFN IFN-λ2/3 and type I and III IFNs restrict enterovirus replication in a virus-specific manner, with type I IFN exhibiting the greatest restriction of E11 and type III IFNs preferentially restricting EV71. Our findings thus define the events associated with EV71 infections in the GI tract, which could lead to the identification of novel therapeutic targets and/or strategies to prevent or treat the pathogenesis and morbidity associated with infections by this virus.

## Results

### Crypt-based monolayer model

Previously, we grew enteroids generated from intestinal crypts isolated from human fetal small intestines cultured in Matrigel and infected these with E11, CVB, and EV71^20^. In this study, we found that EV71 replicated poorly in comparison to other enteroviruses. However, the basis for this low level of infection was unclear. Given that enteroids cultured in Matrigel develop an apical surface facing into the lumen (**Figure 1A**, **Supplemental Figure 1A**), which is not accessible from the culture medium, we theorized that the low levels of EV71 replication in this model might result from the need to infect from the basolateral surface. We therefore determined whether direct culturing of isolated crypts on porous membrane transwell inserts would provide a model to access the apical and basolateral surfaces in an intact monolayer setting. To do this, we isolated intestinal crypts from human fetal small intestines and plated them directly onto T-clear transwell inserts in the presence of factors required to promote stem cell differentiation (R-spondin, Noggin, epidermal growth factor, Wnt3A, and the Rho Kinase inhibitor Y-27632) (**Supplemental Figure 1B**). Similar models have been utilized from crypts isolated from the adult GI tract, which often requires growth as enteroids in Matrigel prior to disruption and subsequent transwell plating (reviewed in ^21^). We found that fetal small intestine-derived crypts plated directly on transwell inserts developed into complete monolayers within 2-3 days post-plating and exhibited distinct apical and basolateral domains that contained distinct intestinal cell types such as mucin-2 (MUC2) positive goblet cells and chromogranin A (CHGA)-positive enteroendocrine cells at the same ratio as crypts cultured in Matrigel (**Figure 1A, Supplemental Figure 1C**). Using RNASeq and RT-qPCR, we found that crypts plated directly in transwell inserts exhibited similar transcriptional profiles (**Supplemental Figure 1D**) and expression of markers of enterocytes (Sucrase-isomaltase (SI), Alkaline Phosphatase (ALPL), goblet cells (MUC2, MUC5AC, MUC13, MUC17), enteroendocrine cells (CHGA), Paneth cells (REG3A), and stem cells (OLFM4), although we did observe significantly lower expression of LGR5 (**Figure 1B, 1C**). In addition to developing a multicellular phenotype, crypt monolayers (hereafter referred to as human intestinal epithelium (HIE)) formed junctional complexes composed of both tight junctions (ZO-1) and adherens junctions (E-cadherin) and exhibited intact barrier function as assessed by high (> approximately 1000 S1 Ω) transepithelial resistance (TER) values (**Figure 1D, 1E**).

**Figure 1.**
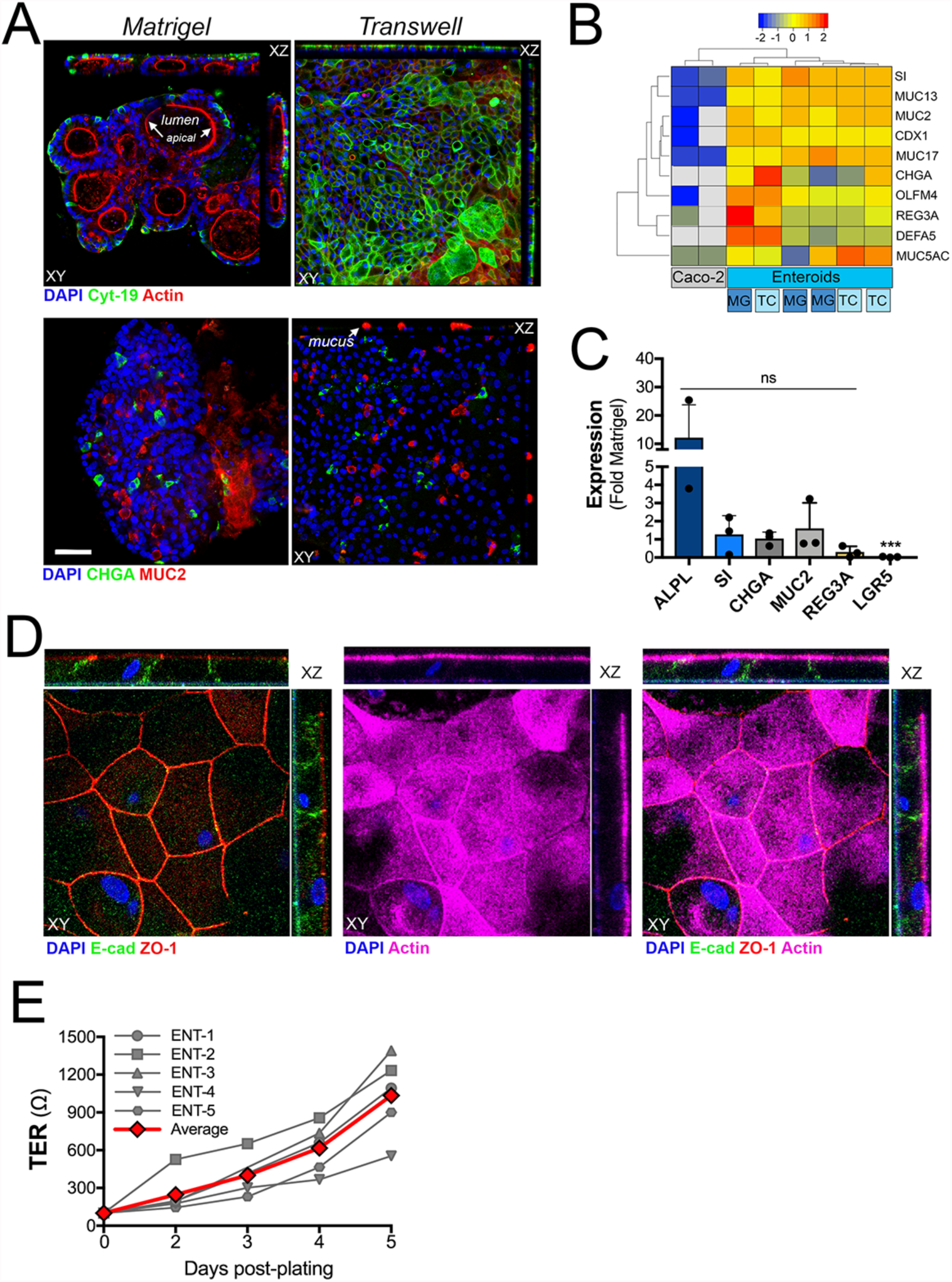
Establishment of human fetal small intestinal-derived monolayer model. **(A),** Confocal micrographs of isolated crypts grown in Matrigel (left) or on transwell T-clear insert (right) for 6 days. Shown is immunofluorescence images from samples immunostained for cytokeratin-19 (an epithelial marker) (green top) and actin (red, top) or chromogranin A (CHGA, an enteroendocrine marker) (green, bottom) and mucin-2 (MUC2, a goblet marker) (red, bottom). In all, DAPI-stained nuclei are shown in blue. At top and right of upper panel are xyz or xzy images obtained by serial sectioning. **(B),** Hierarchical clustering heat map of differential gene expression profiles (based on log2 (RPKM) values) between two independent preparations of Caco-2 cells and three matched independent human enteroid cultures plated in Matrigel (MG) or T-clear transwell inserts (TC) by RNAseq. Key at top (grey indicates no reads mapped). **(C),** RT-qPCR for the indicated markers (alkaline phosphatase (ALPL), sucrase-isomaltase (SI), chromogranin A (CHGA), mucin-2 (MUC2), regenerating islet-derived protein 3 (REG3A), and leucine-rich repeat-containing G-protein coupled receptor 5 (LGR5) in three matched independent human enteroid cultures plated in Matrigel or T-clear transwell inserts. Data are shown as mean ± standard deviation as a fold change from Matrigel-plated enteroids. **(D),** Confocal micrographs of isolated crypts grown on transwell T-clear inserts for 6 days. Shown is immunofluorescence images from samples immunostained for E-cadherin (an adherens junction marker, green), ZO-1 (a tight junction marker, red) and actin (magenta). DAPI-stained nuclei are shown in blue. At top and right of upper panel are xyz or xzy images obtained by serial sectioning. **(E),** Transepithelial resistance (TER, in Ω) values from five independent HIE cultures (ENT-1-5 in grey, two to three transwells were averaged per preparation). Average TER values from all preparations are shown in red.

### EV71 preferentially infects HIE from the apical surface

It is unknown whether enteroviruses exhibit a preferential polarity of binding or infection in primary HIE. To address this, we performed binding and infection assays from either the apical or basolateral surfaces in primary HIE. These studies revealed significant differences in the capacity of E11 and EV71 to bind and infect in a polarized manner. Whereas E11 exhibited an enhanced capacity to infect from the basolateral surface as assessed by the production of vRNA by RT-qPCR at 24hrs post-infection (p.i.), EV71 exhibited a much stronger preference for apical infection (**Figure 2A**). Consistent with this, we found that EV71 preferentially binds to the apical surface of HIE as assessed by a qPCR-based binding assay (**Figure 2B**). To determine whether E11 and EV71 exhibit a polarity of release, we infected HIE with EV71 or E11 from the apical or basolateral surfaces, respectively, and titrated released progeny viral particles from medium isolated from the apical or basolateral compartments. These studies revealed that E11 was released from both the apical and basolateral compartments, although its release was skewed towards the basolateral compartment (**Figure 2C**). In contrast, EV71 was solely released from the apical compartment and no viral particles were detectable in the basolateral compartment (**Figure 2C**).

**Figure 2.**
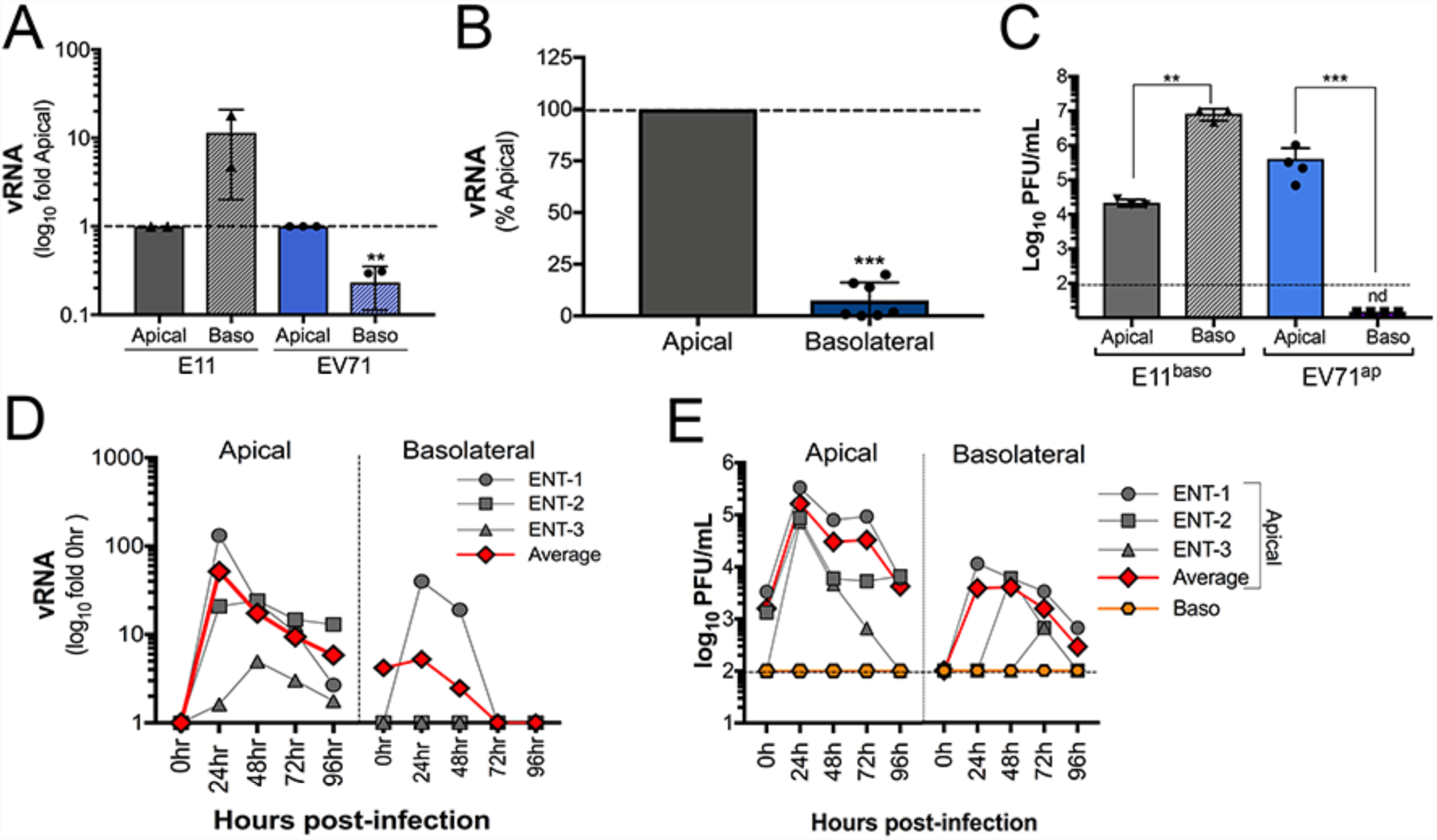
EV71 preferentially infects HIEs from the apical surface. **(A),** E11 and EV71 replication as assessed by the production of vRNA by RT-qPCR when infections were initiated from the apical or basolateral surfaces. Data are shown as fold change from apical infections (log10). Data are from two (E11) or three (EV71 independent HIE cultures. **(B),** Binding efficiency of EV71 when pre-adsorbed to the apical or basolateral surfaces as assessed by RT-qPCR. Data are shown as a percent of apical binding and are from four independent HIE preparations. **(C),** E11 and EV71 replication as assessed by titration of virus from the apical or basolateral compartments when infection was initiated from the apical (EV71) or basolateral (E11) surfaces. Data are from three independent HIE preparations. **(D, E),** Kinetics of neutral-red labeled EV71 growth in three independent HIE preparations at the indicated times. Neutral-red labeled EV71 was pre-adsorbed to the apical or basolateral surfaces for 1hr in the semi-dark, exposed to light at 0hr or at 6hr p.i., and then infection allowed to proceed for indicated hr (24-96hr). Infection was assessed by production of vRNA by RT-qPCR (D) or viral titration (E) from the apical or basolateral (orange) compartments. Note that in (E), no virus was detected in medium isolated from the basolateral compartment. Data are from three independent preparations (ENT-1-3, in grey). Average is shown in red. In (A-C) data are shown as mean ± standard deviation (**P<0.01, ***P<0.001).

We next performed growth curves from HIEs infected with EV71 from either the apical or basolateral surfaces. For these studies, we utilized neutral red (NR)-containing EV71 particles to distinguish between EV71 particles that remained attached to the cell surface from those that were actively replicating. This technique involves labeling of vRNA with NR, a compound that crosslinks the vRNA if exposed to light^22, 23^, thus generating viral particles that are rendered non-infectious when exposed to light. To perform growth curves, NR-EV71 was pre-adsorbed to cells from the apical or basolateral surfaces under semi-dark conditions and exposed to light immediately post-binding (0hr) or following viral entry and genome release (6hr p.i.) and then infected for an additional 24-96hr. NR-EV71 particles that remained at the cell surface would thus be rendered non-infectious at the 6hr light exposure. Using HIEs prepared from three independent human tissues and infected as described, we found that EV71 vRNA production peaked by ∼24h p.i. and then was rapidly reduced by 48-72h p.i., with levels diminishing significantly by 96h p.i. (**Figure 2D**). This trend was specific for apical infection as only a single preparation exhibited any detectable vRNA when infection was initiated from the basolateral surface (**Figure 2D**). In parallel, we collected cell supernatants from the apical or basolateral compartments and measured infectious particle release over a 24-96h period. Consistent with our vRNA data, we found that the levels of infectious EV71 release were highest at 24h p.i., with levels diminishing between 48- 96h p.i. (**Figure 2E**). Of note, even when low levels of infectious EV71 particles were released following infection of the basolateral surface, this release was only detectable in the apical compartment (**Figure 2E**). Taken together, these data show that EV71 exhibits a strong preference to infect HIEs from the apical surface and that infectious particles also exhibit an apical polarity of release.

### EV71 infection of HIE does not alter epithelial barrier function

We showed previously that E11 infection of human enteroids grown in Matrigel induced significant damage to the epithelium, including reorganization of tight junctions^20^. Consistent with this, we found that infection of HIE with E11 from the basolateral surface induced a significant loss of epithelial barrier function, as indicated by the loss of TER values from ∼2000W to ∼200W (**Figure 3A**). In contrast, EV71 infection (from either the apical or basolateral surfaces) had no effect on TER values (**Figure 3A**), even when infection was allowed to proceed for up to 4 days (**Figure 3B**). Likewise, we found that E11 and EV71 also exhibited differences in their impact on epithelial morphology, with E11 infection inducing loss of actin cytoskeletal integrity which was not present in EV71-infected HIE (**Figure 3C**). These data highlight differences amongst enteroviruses on their impact on intestinal epithelial structure and function and show that EV71 infection does not alter epithelial barrier function.

**Figure 3.**
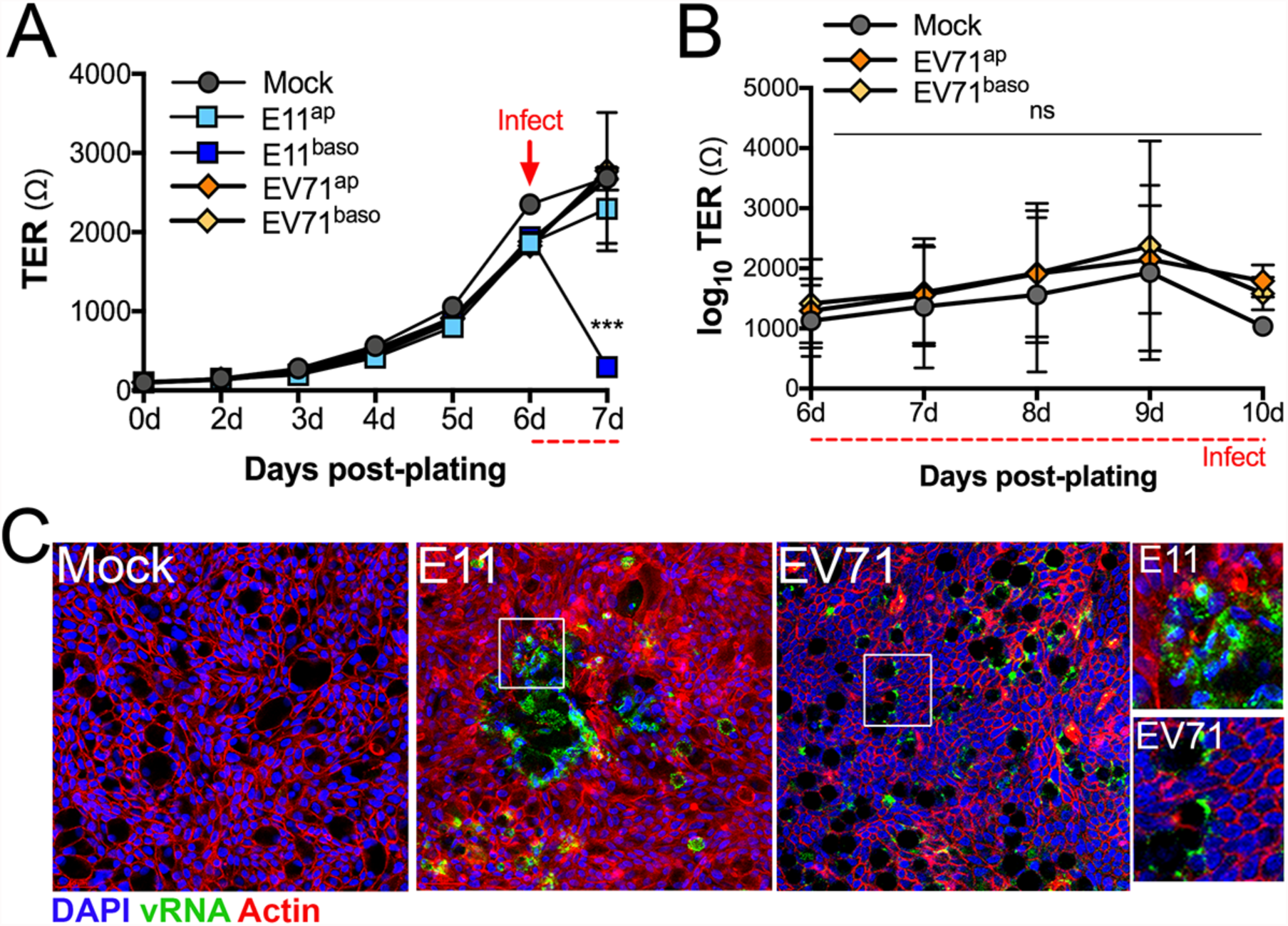
EV71 infection does not induce loss of epithelial barrier integrity. **(A),** TER values at the indicated days post crypt plating in Transwell inserts. At 6d post-plating, Transwells were infected with E11 or EV71 from the apical or basolateral surfaces (red arrow) and TER values were measured 24h post-infection. Data are from a single HIE preparation performed in triplicate and is representative of at least five independent preparations. **(B),** TER values at the indicated days following infection with EV71 from the apical or basolateral surfaces. Data are from four HIE preparation performed in duplicate. **(C),** Confocal microscopy for vRNA (green) or actin (red) in mock infected HIE or HIE infected with E11 from the basolateral surface or EV71 from the apical surface. Images were captured 24h post-infection. Zoomed images from white boxes shown at right. In (A, B) data are shown as mean ± standard deviation (***P<0.001).

### EV71 infects goblet cells

Because we observed differences in the impact of E11 and EV71 infections on epithelial barrier function, we next determined whether these viruses exhibited differences in the specific cell types infected in HIE. We showed previously that E11 preferentially infects enterocytes and can also infect enteroendocrine cells, but is unable to infect goblet cells^20^. To determine if EV71 also exhibits a cell type specificity, we first performed immunofluorescence microscopy for double-stranded vRNA (a replication intermediate) and the virally-encoded capsid protein VP1 in HIE infected with EV71 from the apical surface for 24h (a time when we observed peak levels of replication). These studies revealed colocalization of EV71 vRNA and VP1 to punctate structures in select cells throughout the monolayer (**Figure 4A**). The cells that were positive for EV71 vRNA and VP1 exhibited characteristics of goblet cells, such as a highly polarized nuclear localization and large cytoplasmic space (**Figure 4A**, enlarged panel at right). Indeed, follow up studies confirmed that EV71 vRNA was exclusively localized to mucin-2 (MUC2)-positive goblet cells (**Figure 4B, 4C**). As an additional confirmation for the goblet cell specificity of EV71 infection, we also performed immunofluorescence microscopy for VP1 using an immunostaining technique that distinguishes between VP1 localized on the extracellular surface and VP1 localized intracellularly^24^. These studies confirmed the presence of intracellular VP1 only in cells exhibiting goblet cell morphology (**Figure 4D**). Of note, the primary receptor for EV71, SCARB2^12^, was enriched in goblet cells, where it localized to intracellular vesicles (**Figure 4E**). Consistent with its infection of goblet cells, we also found that EV71 infection of HIE led to significant decreases in the expression of MUC2 as assessed by RT-qPCR, suggesting that infection might alter aspects of goblet cell function (**Figure 4F**). Collectively, these studies show that EV71 specifically infects via the apical surface of HIE and exhibits preferential infection of goblet cells.

**Figure 4.**
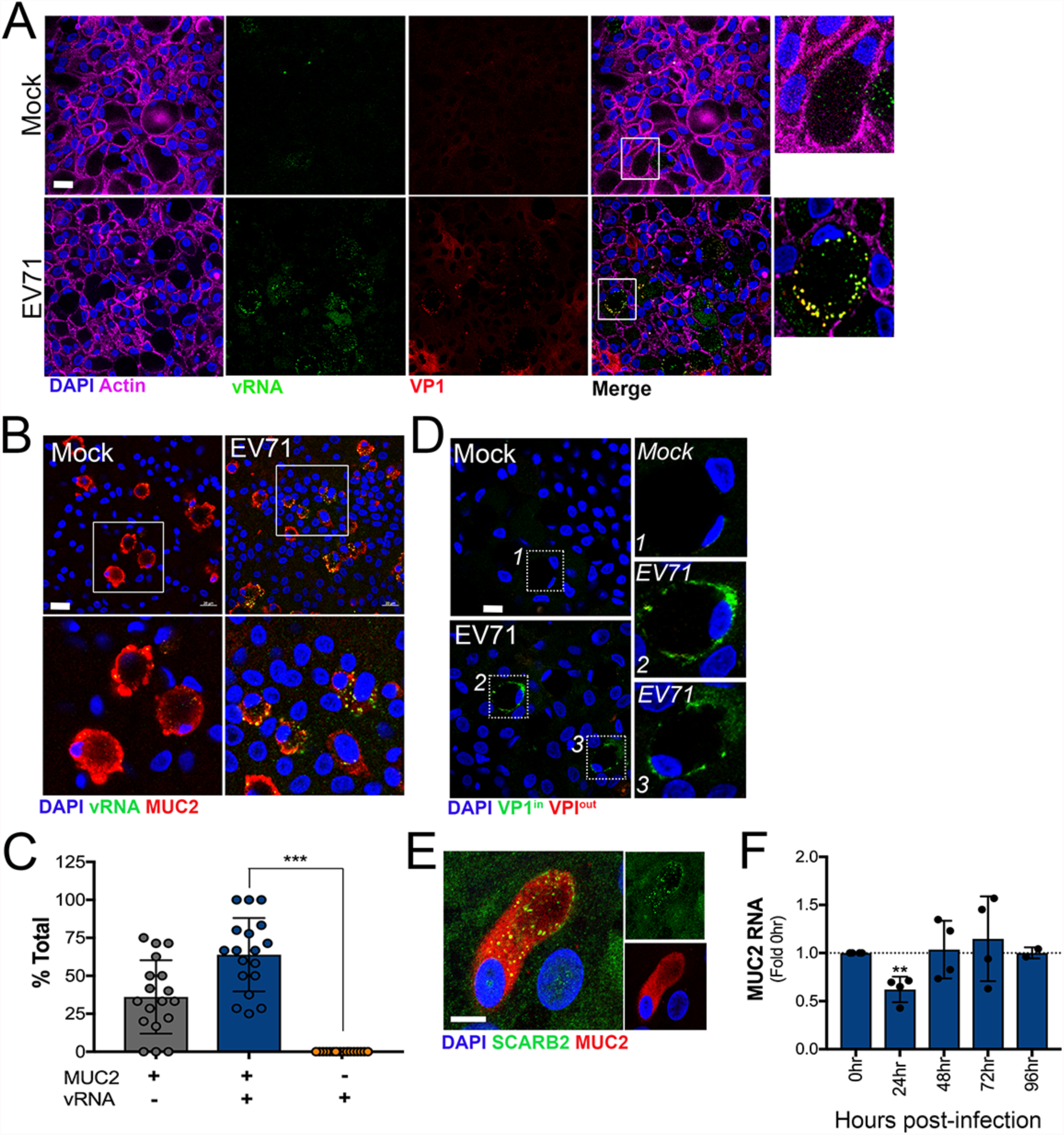
EV71 infects goblet cells. **(A),** Confocal micrographs from mock- or EV71 apically infected HIE immunostained for vRNA (green) and VP1 (red) at 24h post-infection. Zoomed images from areas shown in white boxes shown at right. **(B),** Confocal micrographs from mock- or EV71 apically infected HIE immunostained for vRNA (green) and VP1 (red) at 24h post-infection. Zoomed images from areas shown in white boxes shown at bottom. **(C),** Quantification of the extent of colocalization between vRNA and MUC2-positive or -negative cells as assessed by image analysis. Data were generated from three independent HIE preparations. **(E),** Immunofluorescence microscopy for SCARB2 (green) and MUC2 (red) from HIE grown for 7 days. **(F),** MUC2 expression as assessed by RT-qPCR at the indicated times post-infection (from the apical surface). with neutral-red labeled EV71 exposed to light immediately post-adsorption (0hr) or at 6hr p.i., and then infection allowed to proceed for the indicated time (in hrs). Data are shown as a fold change from HIE exposed to light at 0hr and are from four independent HIE preparations. In (C, D), data are shown as mean ± standard deviation (**P<0.01, ***P<0.001).

### Type III interferons control EV71 infection of HIE

Our EV71 growth curve studies revealed that the peak of EV71 replication was at 24h p.i., with levels of infection declining after this time point (**Figure 2D, 2E**). These data suggest that the host innate immune response to EV71 might suppress viral replication at an early stage in order to control its replication. To determine if this is the case, we performed RT-qPCR analyses for two interferon stimulated genes (ISGs), that we previously showed were induced in HIE in response to E11 infection^20^, in HIE infected with EV71. These studies showed that these ISGs, CXCL10 and IFI44L, were induced by EV71 infection of HIE at 24h p.i., with induction diminishing by 48h-72h p.i. (**Figure 5A**). We next determine whether type I and/or type III IFNs were responsible for this induction of ISGs by performing Luminex singleplex assays for IFNβ, IFNλ-1 or, IFNλ-2/3 (the high degree of sequence homology between these IFNs make them indistinguishable in this assay). We found that EV71 infection of HIE led to the specific induction of type III IFNs, specifically IFNλ-2/3, at both 24h and 48h p.i., with no detectable IFN-λ1 induced and very low levels of IFN-β induced at 24h (**Figure 5B**). Of note, IFNs were present in media collected from the apical chamber following infection and we were unable to detect any IFNs from media collected from the basolateral chamber. Likewise, E11 infection also induced the preferential secretion of type III IFNs, but unlike EV71, but low levels of IFN-λ1 were also produced in response to infection (**Supplemental Figure 2**). These data suggest that type III IFNs, specifically IFN-λ2/3, are induced in response to EV71 infection of HIE.

**Figure 5.**
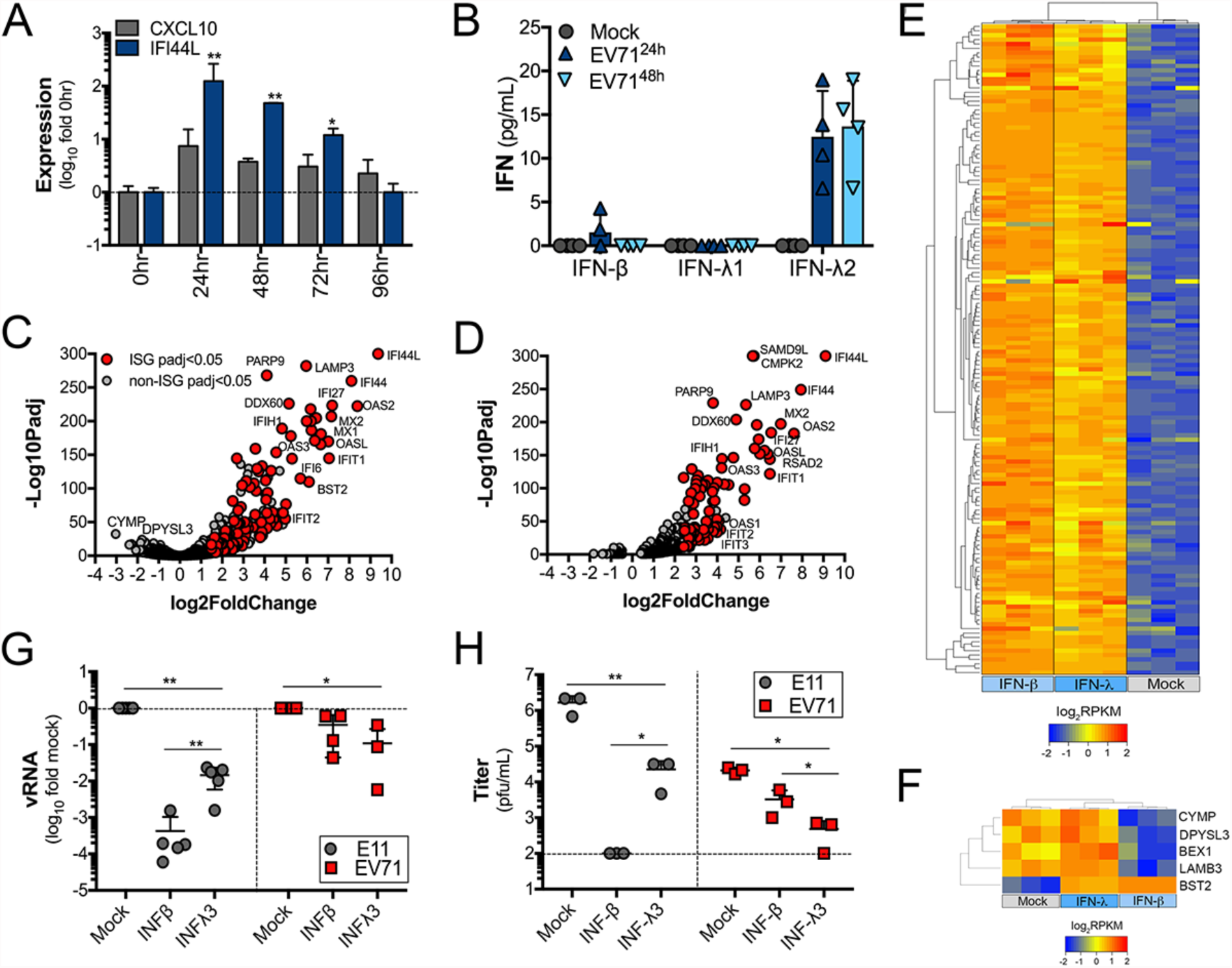
EV71 infection of HIE induces type III IFNs. **(A),** RT-qPCR for two ISGs, CXCL10 and IFI44L, in HIE infected with neutral-red labeled EV71 from the apical surface for the indicated times. Data are from four independent HIE preparations and are shown as a fold change (log10) from cultures exposed to light at 0hr. **(B),** Luminex assays for IFN-β, IFN-λ1, or IFNλ-2/3 from HIE infected with EV71 from the apical surface for 24h or 48h. Data are shown as pg/mL and are from four independent HIE preparations. **(C, D),** Volcano plots of HIE treated with 500ng IFN-β (C) or IFN-λ1 (D) denoting ISGs (red circles) and non-ISGs (grey circles) differentially expressed by treatment (P<0.05). Data are from three independent HIE preparations. **(E),** Hierarchical clustering heatmap (based on log2RPKM values) of canonical ISGs induced by treatment of HIE with IFN-β or IFN-λ1, or mock treated controls. **(G, H),** E11 and EV71 infection from HIE pretreated with 500ng IFN-β or IFN-λ3 for 24h and then infected with E11 or EV71 for 24h. In (G), replication is assessed by vRNA production by RT-qPCR and in (H), viral titration by plaque assay was performed. In (A, B, G, H), data are shown as mean ± standard deviation (*P<0.05, **P<0.01, ***P<0.001).

Next, we determined whether HIE exhibited differences in their ability to respond to exogenous type I and III IFNs and whether these IFNs induced ISGs with differing kinetics, as has been shown in adult enteroids at early time points of exposure^25^. To do this, we first performed RNASeq transcriptional profiling from HIEs treated with recombinant IFN-β or IFN-λ for 24h. Differential expression analysis revealed that fetal-derived HIE potently respond to IFN-β and IFNλ and induce the expression of canonical ISGs to similar levels (**Figure 5C, 5D, 5E**). Moreover, differential expression analysis between IFN-β and IFN-λ treated HIE showed that only five transcripts were differentially regulated by IFN-β, and four of these transcripts were downregulated in response to treatment (**Figure 5F**). A kinetic profiling of the responsiveness of HIE to recombinant IFN-β and IFN-λ confirmed these findings and showed that there were no significant differences in the kinetics by which fetal HIE respond to type I or III IFN treatment (**Supplemental Figure 3**).

Finally, we determined whether E11 and EV71 were differentially controlled by type I or III IFN treatment in a virus- or IFN-specific manner. HIE were pre-treated with recombinant IFN-β or IFN-λ for 24h and then infected with E11 or EV71 from the basolateral or apical surfaces, respectively. We found that whereas E11 was more potently restricted by IFN-β treatment as detected by RT-qPCR for vRNA and viral titration, EV71 was more potently restricted by IFN-λ treatment (**Figure 5G, 5H**). Altogether, these data show that HIE specifically induce type III IFNs in response to EV71 infection, which likely acts to control viral infection at early stages of the viral life cycle.

## Discussion

The events associated with EV71 infection of the human GI tract are largely unknown. Here we show that EV71 preferentially infects HIE from the apical surface where it preferentially replicates in MUC2-positive goblet cells. We also show that unlike E11, an enterovirus that targets enterocytes, EV71 infection of HIE has no impact on epithelial barrier function or cytoskeletal morphology, but infection reduces the expression of MUC2, suggesting that its replication may alter some aspect of goblet cell function. We further show that EV71 infection of HIE induces the type III IFNs IFN-λ2/3, which likely serve to limit EV71 replication. Collectively, these findings provide important insights into the mechanisms by which EV71 and other enteroviruses bypass the GI barrier and point to an important role for type III IFNs in the host response to enterovirus infections within the GI tract.

Our data indicate that enteroviruses exhibit a distinct cell-type specificity by which they infect the human GI tract in a virus-specific manner. Whereas E11 specifically targets enterocytes and also infects enteroendocrine cells^20^, EV71 preferentially infects goblet cells. Although it is possible that EV71 also replicates in other cell types present in HIE at levels that are below the limit of detection of our assays, our data clearly point to an enrichment of EV71 replication in goblet cells. The mechanistic basis for the differential cell type specificity between E11 and EV71 remains unclear, although the cell-type specific expression and localization of viral receptors is likely to play a key role. Although the receptor for E11 is unknown, all EV71 isolates tested to date utilize SCARB2 as a primary receptor^12, 26^. SCARB2, also known as lysosomal integral membrane protein II (LIMPII), is an integral membrane protein that specifically localizes to lysosomes and secretory granules^27^. Indeed, we found that SCARB2 was highly expressed in goblet cells, where it localized to intracellular vesicles. Goblet cells are characterized by the presence of large secretory vesicles that function to transport mucus to the apical surface of the epithelium. The targeting of goblet cells by EV71 for intestinal infection is therefore likely driven at least in part by the enrichment of SCARB2 to secretory vesicles within these cells, which might expose the receptor through apical mucus release. It is also possible that EV71 utilizes other apically-localized attachment factors for its initial binding to the epithelial surface, much like CVB relies on decay accelerating factor (DAF) to attach to the apical surface^28^, before it reaches SCARB2. EV71 has been shown to interact with sialic acid-linked glycans, which might facilitate its initial attachment to the apical surface of the epithelium^29^. However, this binding is unlikely to be a primary determinant for goblet cell infection. The cell-type specific nature of enterovirus infections also suggests that the host response to infection may differ depending on the specific cell types targeted by a given virus. In support of this, our data also point to important differences in the impact of E11 and EV71 infection of epithelial structure and barrier function, which could dramatically impact viral pathogenesis in a virus-specific manner.

Our findings implicate type III IFNs as key contributors in the control of enterovirus infections in the GI tract. These findings are consistent with the work of others who have shown that human rotaviruses^30^^−^^33^, reoviruses^34^, and noroviruses^34^^−^^36^ are also controlled by intestinal-derived type III IFNs. However, unlike other enteric viruses such as rotavirus, which controls the production of type III IFNs during infection through viral antagonism^30^, our findings show that E11 and EV71 infection induce the secretion of type III IFNs at the protein level, suggesting that enteroviruses may lack this mechanism or be less proficient at suppressing this pathway. In cell lines, even those of intestinal lineages, EV71 and other enteroviruses potently antagonize the host innate immune response^37^. This suggests that mechanisms of evasion may differ in primary cells, particularly those isolated from the GI tract. Our data also show that EV71, but not E11, is more potently restricted by type III IFNs than type I IFNs. Similar to E11, rotaviruses are also more sensitive to exogenous treatment with type I IFNs^30^. The mechanistic basis for these differences in sensitivity are unclear, but our data suggest that at least in the fetal GI tract, these differences are unlikely to be the result of differences in the magnitude or kinetics of ISG induction between type I and III IFNs. Instead, these differences may result from differences in the cell type specific nature of enteric virus infections, with rotaviruses^38^ and E11^20^ preferentially infecting enterocytes whereas EV71 targets goblet cells. Dissecting the role of IFNs in the unique cell types of the HIE will likely provide important clues into the differential role that type I and III IFNs might play in the GI tract.

Our studies suggest that enteroviruses have evolved diverse mechanisms to infect distinct cell types in the GI epithelium, which likely impacts many aspects of their pathogenesis, including the role that type III IFNs play in restricting infection and spread. Defining the events associated with EV71 infection in the GI tract could lead to the identification of novel therapeutic targets and/or strategies to prevent or treat the pathogenesis and morbidity associated with infections by this virus.

## Materials and Methods

### Cell culture and human enteroids

Human fetal intestinal crypts were isolated and cultured as described previosuly^20^. Human fetal tissue (< 24 weeks gestation) that resulted from elective terminations were obtained from the University of Pittsburgh Health Sciences Tissue Bank through an honest broker system after approval from the University of Pittsburgh Institutional Review Board and in accordance with the University of Pittsburgh anatomical tissue procurement guidelines. All tissue was genetically normal. Approximately 100 isolated crypts were plated into each well of a 24-well T-clear (0.4µm pore size) transwell insert and were grown in crypt culture media comprised of Advanced DMEM/F12 (Invitrogen) with 20% Hyclone ES Screened Fetal Bovine Serum (Fisher), 1% Penicillin/Streptomycin (Invitrogen), 1% L-glutamine, Gentamycin, 0.2% Amphotericin B, 1% N-acetylcysteine (100mM, Sigma), 1% N-2 supplement (100X, Invitrogen), 2% B27 supplement (50x, Invitrogen), Gibco^®^ HEPES (N-2-hydroxyethylpiperazine-N-2-ethane sulfonic acid, 0.05mM, Invitrogen), ROCK Inhibitor Y-27632 (1mM, 100x, Sigma) and supplemented with the following growth factors 100 ng/ml WNT3a (Fisher), 500 ng/ml R-spondin (R&D), 100 ng/ml Noggin (Peprotech) and 50 ng/ml EGF (Fisher)^39, 40^ for the remainder of the respective experiments, with media changes occurring every 48 hours. Unless otherwise stated, monolayers of HIE were used in studies at six days post-plating.

### Viral infections

Experiments were performed with EV-71 (1095), or E11 (Gregory) that were expanded as described previously^41^. In some cases, experiments were performed with light-sensitive neutral-red viral particles, which was generated as described previously^24^. Briefly, EV71 was propagated in the presence of 10µg/mL of neutral red in the semi-dark and was subsequently purified in semi-dark conditions by ultracentrifugation over a sucrose cushion, as described^41^.

For infections, wells were infected with 10^6 PFU of the indicated virus. Virus was pre-adsorbed to the apical or basolateral surfaces for 1hr at room temperature (basolateral infections were initiated by inverting the transwell inserts). Infections were then initiated by shifting to 37°C and allowed to proceed for the times indicated. For neutral red virus experiments, particles were exposed to light (on a light box) for 20min at 6h p.i. and then infected for the indicated number of hours post-light exposure. In some cases, cells were exposed immediately following adsorption (0hr), which served as a control. E11 and EV71 plaque assays were performed in HeLa cells overlayed with 1.0% or 0.8% agarose respectively; plaques were enumerated following crystal violet staining.

Binding assays were performed by pre-adsorbing 10^6 PFU of the indicated virus to the apical or basolateral surfaces for 60min at room temperature followed by extensive washing with 1x PBS. Following washing, RNA was isolated immediately, and RT-qPCR performed, as described below.

### qPCR and cDNA synthesis

Total RNA was prepared from HIE using the Sigma GenElute total mammalian RNA miniprep kit, according to the protocol of the manufacturer and using the supplementary Sigma DNase digest reagent. RNA was reverse transcribed with the iScript cDNA synthesis kit (Bio-Rad), following the manufacturer’s instructions. 1 μg of total RNA was reversed transcribed in a 20 μL reaction, and subsequently diluted to 100 μL for use. RT-qPCR was performed using the iQ SYBR Green Supermix or iTaq Universal SYBR Green Supermix (Bio-Rad) on a CFX96 Touch Real-Time PCR Detection System (Bio-Rad). Gene expression was determined based on a Δ*CQ* method, normalized to human actin. Primer sequences can be found in Supplemental Table 1.

### RNASeq

Total RNA was extracted as described above. RNA quality was assessed by NanoDrop and an Agilent bioanalyzer and 1µg was used for library preparation using the TruSeq Stranded mRNA Library Preparation kit (Illumina) per the manufacturer’s instructions. Sequencing was performed on an Illumina Nextseq 500. RNAseq FASTQ data were processed and mapped to the human reference genome (hg38) using CLC Genomics Workbench 11 (Qiagen). CLC Genomics was also was used to determine differentially expressed genes at a significance cutoff of p<0.05, unless otherwise stated. Hierarchical gene expression clustering was performed using Cluster 3.0, using average linkage clustering of genes centered by their mean RPKM values. Heat maps (based on log2(RPKM) values) were generated in Heatmapper^42^. Analysis of the transcriptional profile of Caco-2 cells were based on previously published datasets^43^ which were deposited in sequence read archives (SRA) SRP065330. Files from HIE used in the current study were deposited in SRA.

### Immunofluorescence microscopy

Monolayers grown on transwell inserts were washed with PBS and fixed with 4% paraformaldehyde at room temperature, followed by 0.25% Triton X-100 to permeabilize cell membranes for 30min at room temperature. Cultures were incubated with primary antibodies for 1 hour at room temperature, washed, and then incubated for 30 minutes at room temperature with Alexa-Fluor-conjugated secondary antibodies (Invitrogen). Slides were washed and mounted with Vectashield (Vector Laboratories) containing 4′,6-diamidino-2-phenylindole (DAPI). The following antibodies or reagents were used—recombinant anti-dsRNA antibody (provided by Abraham Brass, University of Massachusetts and described previously^44^), Mucin-2 (H-300, Santa Cruz Biotechnology), Lysozyme C (E-5, Santa Cruz Biotechnology), E-cadherin (ECCD-2, Invitrogen), ZO-1 (ZMD.436, Invitrogen), Cytokeratin-19 (EP1580Y, Abcam), VP1 (NCL-ENTERO, Leica), and SCARB2 (EPR12081, Abcam) and Alexa Fluor 594 or 633 conjugated Phalloidin (Invitrogen). Images were captured using a Zeiss LSM 710 inverted laser scanning confocal microscope or with a Leica SP8X tandem scanning confocal microscope with white light laser and contrast adjusted in Photoshop. Image analysis was performed using Fiji. MUC2 and VP1 positive cells were counted using the ImageJ Cell Counter plugin.

### Recombinant IFN treatments

HIE monolayers were treated with 100-500ng of recombinant IFN-β, IFN-λ1 or IFN-λ3 (R&D Systems; 1598-IL-025, 5259-IL-025, 8499-IF-010) added to both the apical and basolateral compartments for ∼20h prior to initiating infections, as described above.

### Luminex assays

Luminex profiling was performed using the Human Bio-Plex Pro Inflammation Panel 1 IFN-β, IL-29, and IL28A sets (Bio-Rad) according to the manufacturer’s protocol using the laboratory multianalyte profiling system (LabMAP^TM^) system developed by Luminex Corporation (Austin, TX).

### Statistics

All statistical analysis was performed using GraphPad Prism. Experiments were performed at least three times from independent intestines (a total of 29 intestines were used in this study) as indicated in the figure legends or as detailed. Data are presented as mean ± standard deviation. Except were specified, a Student’s t-test was used to determine statistical significance. P values of < 0.05 were considered statistically significant, with specific P-values noted in the figure legends.

## Acknowledgements

We thank Kevin McHugh (Children’s Hospital of Pittsburgh) for assistance with Luminex assays, William Horne (Children’s Hospital of Pittsburgh) for assistance with RNASeq, Abraham Brass (University of Massachusetts) for providing anti-dsRNA antibody, and Coyne Drummond (University of Pittsburgh) for technical assistance. This project was supported by NIH R01- AI081759 (C.B.C.) and a Burroughs Wellcome Investigators in the Pathogenesis of Infectious Disease Award (C.B.C), and the Children’s Hospital of Pittsburgh of the UPMC Health System (C.B.C.). The authors would also like to acknowledge the Tissue and Research Pathology Services/Health Sciences Tissue Bank, which receives funding from P30CA047904.

---

**Supplemental Figure 1.**
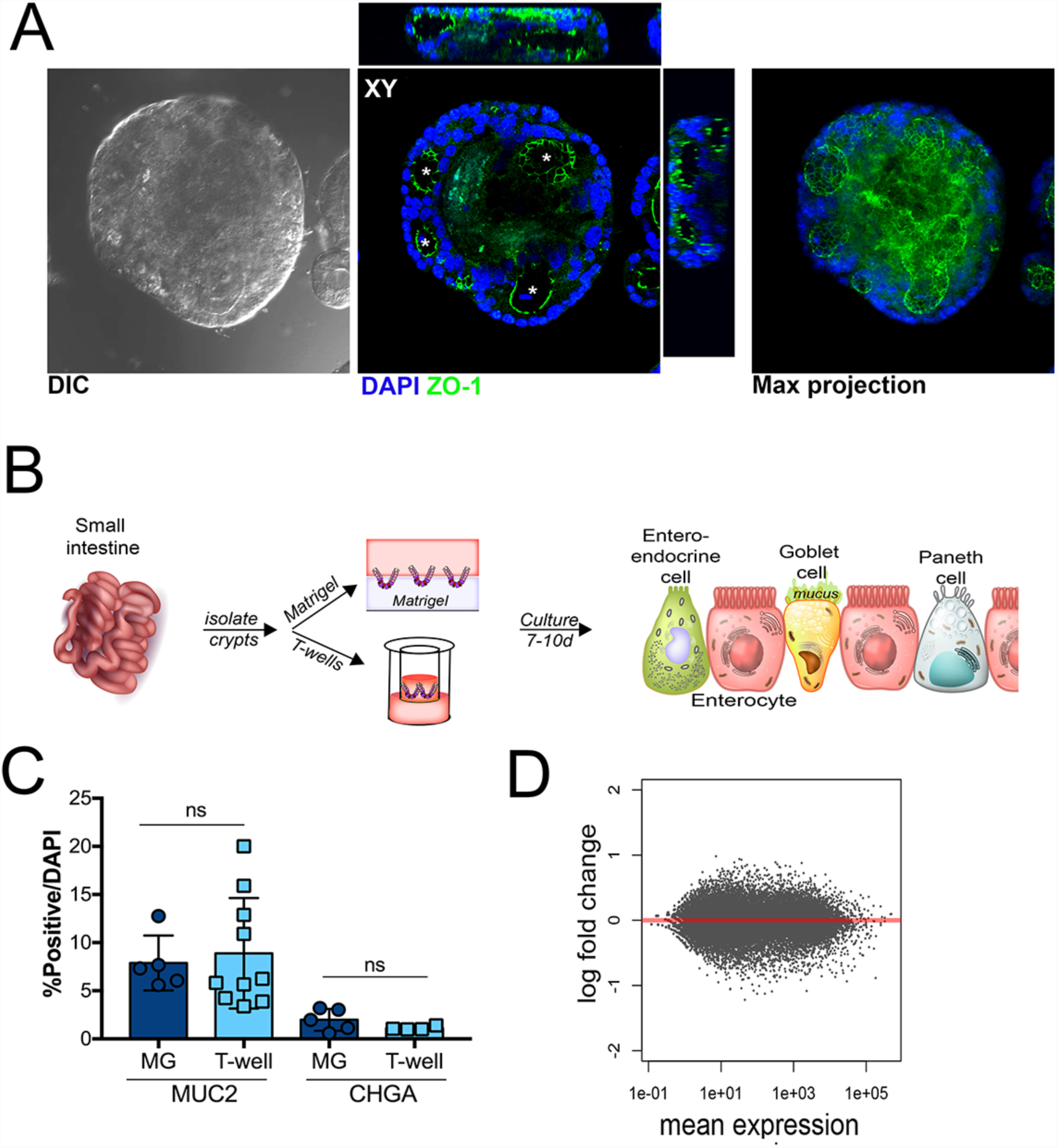
**(A),** Confocal micrograph of enteroid grown in Matrigel for 7 days and immunostained for ZO-1 (in green). DAPI-stained nuclei are shown in blue. At middle, xzy and xyz cross-sections are shown at top and right. At right, maximum projection is shown. **(B),** Schematic of the isolation and culturing of isolated crypts in Matrigel or on Transwells. **(C),** Quantification of the numbers of MUC2 and CHGA positive cells from crypts isolated and grown in Matrigel (MG) or on Transwell inserts (T-well) (normalized to DAPI). Data are shown as mean ± standard deviation and were calculated from three independent preparations. **(D),** MA plot generated in R following DeSeq2 analysis demonstrating the differential expression of transcripts between crypts cultured in Matrigel or on Transwell inserts. Data are plotted as log2 fold changes (y-axis) and mean expression (x-axis). Grey denotes transcripts not differentially expressed ( p<0.05).

**Supplemental Figure 2.**
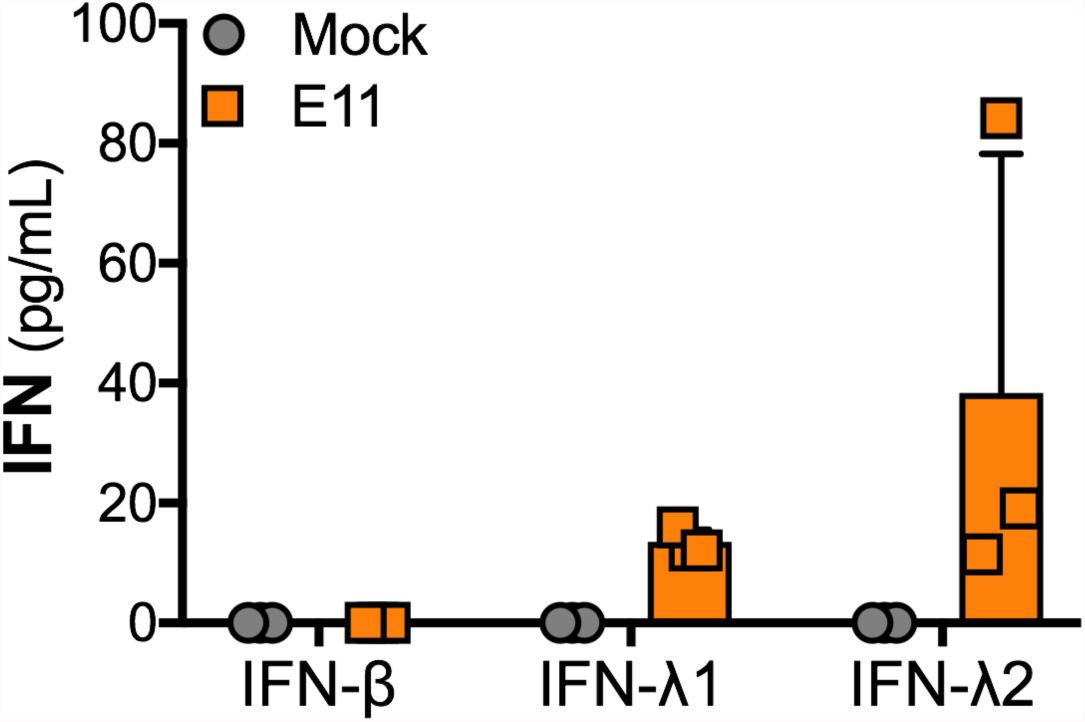
Luminex assays for IFN-β, IFN-λ1, or IFNλ-2/3 from enteroids infected with E11 for 24h. Data are shown as pg/mL and are from three independent preparations

**Supplemental Figure 3.**
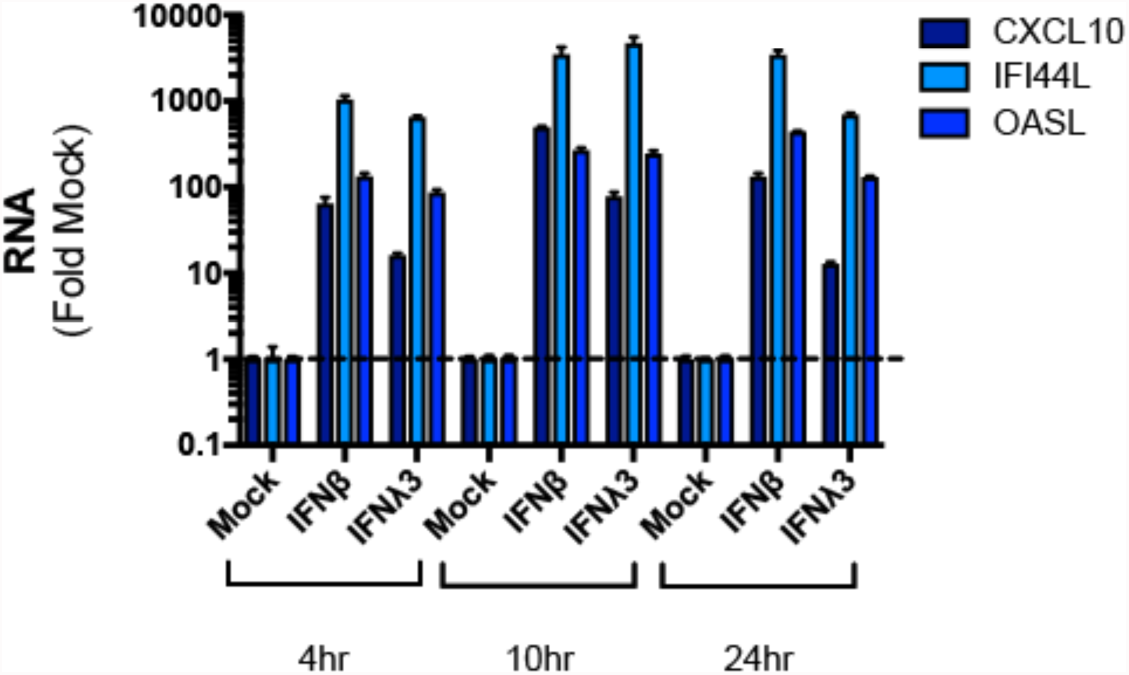
RT-qPCR for the ISGs CXCL10, IFI44L, and OASL from human enteroids treated with 100ng/mL of IFN-β or IFN-λ3 for the indicated times. Data are shown as mean ± standard deviation normalized to mock-treated controls.

**Supplemental Table 1.**
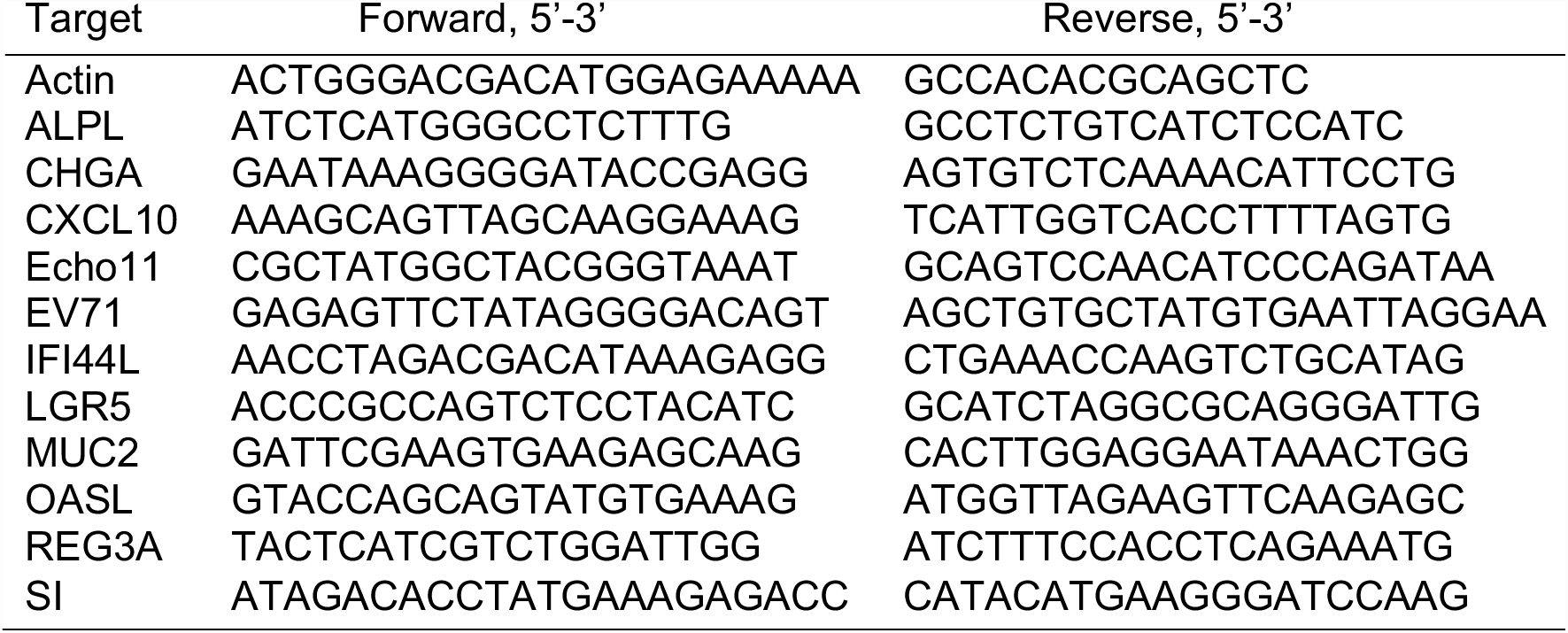
Primers used in this study

## Literature Cited

1. Yip CC, Lau SK, Woo PC, Yuen KY. Human enterovirus 71 epidemics: what’s next? Emerg Health Threats J. 2013;6:19780. Epub 2013/10/15. doi: 10.3402/ehtj.v6i0.19780. PubMed PMID: 24119538; PMCID: PMC3772321.

2. Schmidt NJ, Lennette EH, Ho HH. An apparently new enterovirus isolated from patients with disease of the central nervous system. J Infect Dis. 1974;129(3):304–9. Epub 1974/03/01. PubMed PMID: 4361245.

3. Xing W, Liao Q, Viboud C, Zhang J, Sun J, Wu JT, Chang Z, Liu F, Fang VJ, Zheng Y, Cowling BJ, Varma JK, Farrar JJ, Leung GM, Yu H. Hand, foot, and mouth disease in China, 2008-12: an epidemiological study. Lancet Infect Dis. 2014;14(4):308–18. Epub 2014/02/04. doi: 10.1016/S1473-3099(13)70342-6. PubMed PMID: 24485991; PMCID: PMC4035015.

4. Wang SM, Liu CC, Tseng HW, Wang JR, Huang CC, Chen YJ, Yang YJ, Lin SJ, Yeh TF. Clinical spectrum of enterovirus 71 infection in children in southern Taiwan, with an emphasis on neurological complications. Clin Infect Dis. 1999;29(1):184–90. Epub 1999/08/05. doi: 10.1086/520149. PubMed PMID: 10433583.

5. Chang LY, Lin TY, Hsu KH, Huang YC, Lin KL, Hsueh C, Shih SR, Ning HC, Hwang MS, Wang HS, Lee CY. Clinical features and risk factors of pulmonary oedema after enterovirus-71- related hand, foot, and mouth disease. Lancet. 1999;354(9191):1682–6. Epub 1999/11/24. doi: 10.1016/S0140-6736(99)04434-7. PubMed PMID: 10568570.

6. Liu CC, Tseng HW, Wang SM, Wang JR, Su IJ. An outbreak of enterovirus 71 infection in Taiwan, 1998: epidemiologic and clinical manifestations. J Clin Virol. 2000;17(1):23–30. Epub 2000/05/18. PubMed PMID: 10814935.

7. Chen YC, Yu CK, Wang YF, Liu CC, Su IJ, Lei HY. A murine oral enterovirus 71 infection model with central nervous system involvement. J Gen Virol. 2004;85(Pt 1):69–77. Epub 2004/01/14. doi: 10.1099/vir.0.19423-0. PubMed PMID: 14718621.

8. Wang YF, Chou CT, Lei HY, Liu CC, Wang SM, Yan JJ, Su IJ, Wang JR, Yeh TM, Chen SH, Yu CK. A mouse-adapted enterovirus 71 strain causes neurological disease in mice after oral infection. J Virol. 2004;78(15):7916–24. Epub 2004/07/16. doi: 10.1128/JVI.78.15.7916- 7924.2004. PubMed PMID: 15254164; PMCID: PMC446098.

9. Zhang Y, Cui W, Liu L, Wang J, Zhao H, Liao Y, Na R, Dong C, Wang L, Xie Z, Gao J, Cui P, Zhang X, Li Q. Pathogenesis study of enterovirus 71 infection in rhesus monkeys. Lab Invest. 2011;91(9):1337–50. Epub 2011/05/11. doi: 10.1038/labinvest.2011.82. PubMed PMID: 21555996.

10. Khong WX, Yan B, Yeo H, Tan EL, Lee JJ, Ng JK, Chow VT, Alonso S. A non-mouse-adapted enterovirus 71 (EV71) strain exhibits neurotropism, causing neurological manifestations in a novel mouse model of EV71 infection. J Virol. 2012;86(4):2121–31. Epub 2011/12/02. doi: 10.1128/JVI.06103-11. PubMed PMID: 22130542; PMCID: PMC3302383.

11. Fujii K, Nagata N, Sato Y, Ong KC, Wong KT, Yamayoshi S, Shimanuki M, Shitara H, Taya C, Koike S. Transgenic mouse model for the study of enterovirus 71 neuropathogenesis. Proc Natl Acad Sci U S A. 2013;110(36):14753–8. Epub 2013/08/21. doi: 10.1073/pnas.1217563110. PubMed PMID: 23959904; PMCID: PMC3767555.

12. Yamayoshi S, Yamashita Y, Li J, Hanagata N, Minowa T, Takemura T, Koike S. Scavenger receptor B2 is a cellular receptor for enterovirus 71. Nat Med. 2009;15(7):798–801. Epub 2009/06/23. doi: 10.1038/nm.1992. PubMed PMID: 19543282.

13. Nagata N, Shimizu H, Ami Y, Tano Y, Harashima A, Suzaki Y, Sato Y, Miyamura T, Sata T, Iwasaki T. Pyramidal and extrapyramidal involvement in experimental infection of cynomolgus monkeys with enterovirus 71. J Med Virol. 2002;67(2):207–16. Epub 2002/05/07. doi: 10.1002/jmv.2209. PubMed PMID: 11992581.

14. Nagata N, Iwasaki T, Ami Y, Tano Y, Harashima A, Suzaki Y, Sato Y, Hasegawa H, Sata T, Miyamura T, Shimizu H. Differential localization of neurons susceptible to enterovirus 71 and poliovirus type 1 in the central nervous system of cynomolgus monkeys after intravenous inoculation. J Gen Virol. 2004;85(Pt 10):2981–9. Epub 2004/09/28. doi: 10.1099/vir.0.79883-0. PubMed PMID: 15448361.

15. Sato T, van Es JH, Snippert HJ, Stange DE, Vries RG, van den Born M, Barker N, Shroyer NF, van de Wetering M, Clevers H. Paneth cells constitute the niche for Lgr5 stem cells in intestinal crypts. Nature. 2011;469(7330):415–8. Epub 2010/11/30. doi: 10.1038/nature09637. PubMed PMID: 21113151; PMCID: PMC3547360.

16. Kretzschmar K, Clevers H. Organoids: Modeling Development and the Stem Cell Niche in a Dish. Dev Cell. 2016;38(6):590–600. doi: 10.1016/j.devcel.2016.08.014. PubMed PMID: 27676432.

17. In JG, Foulke-Abel J, Estes MK, Zachos NC, Kovbasnjuk O, Donowitz M. Human mini-guts: new insights into intestinal physiology and host-pathogen interactions. Nat Rev Gastroenterol Hepatol. 2016. doi: 10.1038/nrgastro.2016.142. PubMed PMID: 27677718.

18. Date S, Sato T. Mini-gut organoids: reconstitution of the stem cell niche. Annu Rev Cell Dev Biol. 2015;31:269–89. doi: 10.1146/annurev-cellbio-100814-125218. PubMed PMID: 26436704.

19. Lanik WE, Mara MA, Mihi B, Coyne CB, Good M. Stem Cell-Derived Models of Viral Infections in the Gastrointestinal Tract. Viruses. 2018;10(3). Epub 2018/03/15. doi: 10.3390/v10030124. PubMed PMID: 29534451; PMCID: PMC5869517.

20. Drummond CG, Bolock AM, Ma C, Luke CJ, Good M, Coyne CB. Enteroviruses infect human enteroids and induce antiviral signaling in a cell lineage-specific manner. Proc Natl Acad Sci U S A. 2017;114(7):1672–7. Epub 2017/02/01. doi: 10.1073/pnas.1617363114. PubMed PMID: 28137842; PMCID: PMC5320971.

21. In JG, Foulke-Abel J, Estes MK, Zachos NC, Kovbasnjuk O, Donowitz M. Human mini-guts: new insights into intestinal physiology and host-pathogen interactions. Nat Rev Gastroenterol Hepatol. 2016;13(11):633–42. Epub 2016/10/26. doi: 10.1038/nrgastro.2016.142. PubMed PMID: 27677718; PMCID: PMC5079760.

22. Crowther D, Melnick JL. The incorporation of neutral red and acridine orange into developing poliovirus particles making them photosensitive. Virology. 1961;14:11–21. PubMed PMID: 13696675.

23. Brandenburg B, Lee LY, Lakadamyali M, Rust MJ, Zhuang X, Hogle JM. Imaging poliovirus entry in live cells. PLoS Biol. 2007;5(7):e183. doi: 10.1371/journal.pbio.0050183.

24. Delorme-Axford E, Sadovsky Y, Coyne CB. Lipid raft- and SRC family kinase-dependent entry of coxsackievirus B into human placental trophoblasts. J Virol. 2013;87(15):8569–81. Epub 2013/05/31. doi: 10.1128/JVI.00708-13. PubMed PMID: 23720726; PMCID: PMC3719791.

25. Pervolaraki K, Stanifer ML, Munchau S, Renn LA, Albrecht D, Kurzhals S, Senis E, Grimm D, Schroder-Braunstein J, Rabin RL, Boulant S. Type I and Type III Interferons Display Different Dependency on Mitogen-Activated Protein Kinases to Mount an Antiviral State in the Human Gut. Front Immunol. 2017;8:459. Epub 2017/05/10. doi: 10.3389/fimmu.2017.00459. PubMed PMID: 28484457; PMCID: PMC5399069.

26. Yamayoshi S, Iizuka S, Yamashita T, Minagawa H, Mizuta K, Okamoto M, Nishimura H, Sanjoh K, Katsushima N, Itagaki T, Nagai Y, Fujii K, Koike S. Human SCARB2-dependent infection by coxsackievirus A7, A14, and A16 and enterovirus 71. J Virol. 2012;86(10):5686–96. Epub 2012/03/23. doi: 10.1128/JVI.00020-12. PubMed PMID: 22438546; PMCID: PMC3347270.

27. Vega MA, Segui-Real B, Garcia JA, Cales C, Rodriguez F, Vanderkerckhove J, Sandoval IV. Cloning, sequencing, and expression of a cDNA encoding rat LIMP II, a novel 74- kDa lysosomal membrane protein related to the surface adhesion protein CD36. J Biol Chem. 1991;266(25):16818–24. Epub 1991/09/05. PubMed PMID: 1715871.

28. Shieh JT, Bergelson JM. Interaction with decay-accelerating factor facilitates coxsackievirus B infection of polarized epithelial cells. J Virol. 2002;76(18):9474–80. Epub 2002/08/21. PubMed PMID: 12186929; PMCID: PMC136423.

29. Yang B, Chuang H, Yang KD. Sialylated glycans as receptor and inhibitor of enterovirus 71 infection to DLD-1 intestinal cells. Virol J. 2009;6:141. Epub 2009/09/16. doi: 10.1186/1743- 422X-6-141. PubMed PMID: 19751532; PMCID: PMC2751754.

30. Saxena K, Simon LM, Zeng XL, Blutt SE, Crawford SE, Sastri NP, Karandikar UC, Ajami NJ, Zachos NC, Kovbasnjuk O, Donowitz M, Conner ME, Shaw CA, Estes MK. A paradox of transcriptional and functional innate interferon responses of human intestinal enteroids to enteric virus infection. Proc Natl Acad Sci U S A. 2017;114(4):E570–E9. Epub 2017/01/11. doi: 10.1073/pnas.1615422114. PubMed PMID: 28069942; PMCID: PMC5278484.

31. Hernandez PP, Mahlakoiv T, Yang I, Schwierzeck V, Nguyen N, Guendel F, Gronke K, Ryffel B, Hoelscher C, Dumoutier L, Renauld JC, Suerbaum S, Staeheli P, Diefenbach A. Interferon-lambda and interleukin 22 act synergistically for the induction of interferon-stimulated genes and control of rotavirus infection. Nat Immunol. 2015;16(7):698–707. Epub 2015/05/26. doi: 10.1038/ni.3180. PubMed PMID: 26006013; PMCID: PMC4589158.

32. Pott J, Mahlakoiv T, Mordstein M, Duerr CU, Michiels T, Stockinger S, Staeheli P, Hornef MW. IFN-lambda determines the intestinal epithelial antiviral host defense. Proc Natl Acad Sci U S A. 2011;108(19):7944–9. Epub 2011/04/27. doi: 10.1073/pnas.1100552108. PubMed PMID: 21518880; PMCID: PMC3093475.

33. Lin JD, Feng N, Sen A, Balan M, Tseng HC, McElrath C, Smirnov SV, Peng J, Yasukawa LL, Durbin RK, Durbin JE, Greenberg HB, Kotenko SV. Distinct Roles of Type I and Type III Interferons in Intestinal Immunity to Homologous and Heterologous Rotavirus Infections. PLoS Pathog. 2016;12(4):e1005600. Epub 2016/04/30. doi: 10.1371/journal.ppat.1005600. PubMed PMID: 27128797; PMCID: PMC4851417.

34. Baldridge MT, Lee S, Brown JJ, McAllister N, Urbanek K, Dermody TS, Nice TJ, Virgin HW. Expression of Ifnlr1 on Intestinal Epithelial Cells Is Critical to the Antiviral Effects of Interferon Lambda against Norovirus and Reovirus. J Virol. 2017;91(7). Epub 2017/01/13. doi: 10.1128/JVI.02079-16. PubMed PMID: 28077655; PMCID: PMC5355594.

35. Nice TJ, Baldridge MT, McCune BT, Norman JM, Lazear HM, Artyomov M, Diamond MS, Virgin HW. Interferon-lambda cures persistent murine norovirus infection in the absence of adaptive immunity. Science. 2015;347(6219):269–73. Epub 2014/11/29. doi: 10.1126/science.1258100. PubMed PMID: 25431489; PMCID: PMC4398891.

36. Baldridge MT, Nice TJ, McCune BT, Yokoyama CC, Kambal A, Wheadon M, Diamond MS, Ivanova Y, Artyomov M, Virgin HW. Commensal microbes and interferon-lambda determine persistence of enteric murine norovirus infection. Science. 2015;347(6219):266–9. Epub 2014/11/29. doi: 10.1126/science. 1258025. PubMed PMID: 25431490; PMCID: PMC4409937.

37. Harris KG, Coyne CB. Enter at your own risk: how enteroviruses navigate the dangerous world of pattern recognition receptor signaling. Cytokine. 2013;63(3):230–6. Epub 2013/06/15. doi: 10.1016/j.cyto.2013.05.007. PubMed PMID: 23764548; PMCID: PMC3987772.

38. Saxena K, Blutt SE, Ettayebi K, Zeng XL, Broughman JR, Crawford SE, Karandikar UC, Sastri NP, Conner ME, Opekun AR, Graham DY, Qureshi W, Sherman V, Foulke-Abel J, In J, Kovbasnjuk O, Zachos NC, Donowitz M, Estes MK. Human Intestinal Enteroids: a New Model To Study Human Rotavirus Infection, Host Restriction, and Pathophysiology. J Virol. 2016;90(1):43–56. Epub 2015/10/09. doi: 10.1128/JVI.01930-15. PubMed PMID: 26446608; PMCID: PMC4702582.

39. Shaffiey SA, Jia H, Keane T, Costello C, Wasserman D, Quidgley M, Dziki J, Badylak S, Sodhi CP, March JC, Hackam DJ. Intestinal stem cell growth and differentiation on a tubular scaffold with evaluation in small and large animals. Regen Med. 2016;11(1):45–61. doi: 10.2217/rme.15.70. PubMed PMID: 26395928; PMCID: PMC4891976.

40. Egan CE, Sodhi CP, Good M, Lin J, Jia H, Yamaguchi Y, Lu P, Ma C, Branca MF, Weyandt S, Fulton WB, Nino DF, Prindle T, Jr., Ozolek JA, Hackam DJ. Toll-like receptor 4- mediated lymphocyte influx induces neonatal necrotizing enterocolitis. J Clin Invest. 2016;126(3):495–508. doi: 10.1172/JCI83356. PubMed PMID: 26690704; PMCID: PMC4731173.

41. Morosky S, Lennemann NJ, Coyne CB. BPIFB6 Regulates Secretory Pathway Trafficking and Enterovirus Replication. J Virol. 2016;90(10):5098–107. doi: 10.1128/JVI.00170- 16. PubMed PMID: 26962226; PMCID: PMC4859712.

42. Babicki S, Arndt D, Marcu A, Liang Y, Grant JR, Maciejewski A, Wishart DS. Heatmapper: web-enabled heat mapping for all. Nucleic Acids Res. 2016;44(W1):W147–53. Epub 2016/05/18. doi: 10.1093/nar/gkw419. PubMed PMID: 27190236; PMCID: PMC4987948.

43. Drummond CG, Nickerson CA, Coyne CB. A Three-Dimensional Cell Culture Model To Study Enterovirus Infection of Polarized Intestinal Epithelial Cells. mSphere. 2016;1(1). doi: 10.1128/mSphere.00030-15. PubMed PMID: 27303677; PMCID: PMC4863623.

44. Savidis G, Perreira JM, Portmann JM, Meraner P, Guo Z, Green S, Brass AL. The IFITMs Inhibit Zika Virus Replication. Cell Rep. 2016;15(11):2323–30. doi: 10.1016/j.celrep.2016.05.074. PubMed PMID: 27268505.

